# emeraLD: Rapid Linkage Disequilibrium Estimation with Massive Data Sets

**DOI:** 10.1101/301366

**Authors:** Corbin Quick, Christian Fuchsberger, Daniel Taliun, Gonçalo Abecasis, Michael Boehnke, Hyun Min Kang

## Abstract

**Summary:** Estimating linkage disequilibrium (LD) is essential for a wide range of summary statistics-based association methods for genome-wide association studies (GWAS). Large genetic data sets, e.g. the TOPMed WGS project and UK Biobank, enable more accurate and comprehensive LD estimates, but increase the computational burden of LD estimation. Here, we describe emeraLD (Efficient Methods for Estimation and Random Access of LD), a computational tool that leverages sparsity and haplotype structure to estimate LD orders of magnitude faster than existing tools.

**Availability and Implementation:** emeraLD is implemented in C++, and is open source under GPLv3. Source code, documentation, an R interface, and utilities for analysis of summary statistics are freely available at http://github.com/statgen/emeraLD

**Supplementary information:** Supplementary data are available at *Bioinformatics* online.

## 1 Introduction

Linkage disequilibrium (LD) – pairwise association between alleles at different genetic variants – is of fundamental interest in population genetics as a vestige of natural selection and demographic history, and is essential for a wide range of analyses from summary statistics in genome-wide association studies (GWAS). Motivated by restrictive data sharing policies and logistical constraints, a variety of methods have been developed for analysis of GWAS summary statistics (single-variant association statistics) rather than individual-level data. For example, summary statistics-based methods have been developed for fine-mapping (Benner *et al.*, 2016), conditional association (Yang *et al.*, 2012), gene-based association (Bakshi *et al.*, 2016; Barbeira *et al.*, 2016; Lamparter *et al.*, 2016), heritability estimation (Bakshi *et al.*, 2016), and functional enrichment analysis (Finucane *et al.*, 2015; Lamparter *et al.*, 2016). These methods generally rely on LD estimates from an external data set, which are ideally calculated on-the-fly rather than precomputed and stored due to prohibitive storage costs. For example, the 1000 Genomes Project Phase 3 panel includes over 35M shared variants (1000 Genomes Project Consortium, 2015), which corresponds to over 4 × 10^11^ pairwise LD coefficients within 1 Mbp windows genome-wide.

### 1.1 Existing Tools to Estimate LD

Existing tools to estimate LD generally scale linearly with sample size, prompting a need for more efficient methods for large data sets. PLINK is a widely used software toolkit for analyzing genetic data, and is among the most computationally efficient tools for estimating LD (Purcell *et al.*, 2007; Purcell and Chang, 2016). PLINK’s BED genotype data format allows efficient querying and data processing, but demands prohibitive storage space for large sample sizes and large numbers of markers (e.g., 7.6TB for the TOPMed Whole Genome Sequencing Project, which includes >60K individuals). VCFtools is another widely used software toolkit for manipulating and analyzing genetic data in the Variant Call Format (VCF) (Danecek *et al.*, 2011). Compressed VCF files (VCF.gz) require far less storage space than BED files (e.g., >30× less storage space for the TOPMed WGS Project), and permit random access of genomic regions through block-compression and Tabix indexing (Danecek *et al.*, 2011; Li, 2011). VCFtools provides utilities to estimate LD from VCF files, but is computationally burdensome for large data sets. M3VCF format uses a compact haplotype representation that requires far less storage than genotype formats (Das *et al.*, 2016). m3vcftools provides efficient utilities for estimating LD with M3VCF format, but is substantially slower than PLINK with BED file input.

## 2 Methods

### 2.1 LD Statistics

Three common measures of LD are the LD coefficient *D* (the covariance of genotypes), the standardized LD coefficient *D′* (*D* divided by its maximum value given allele frequencies), and the Pearson correlation *r* or its square (Gabriel *et al.*, 2002). Each of these statistics can be written as a function of allele frequency estimates, sample size, and dot product of genotype vectors. Importantly, only the dot product must be calculated for each pair of variants to calculate LD, since allele frequencies and haplotype counts can be precomputed when processing genotype data.

### 2.2 Computational Approach

We tailored our computational approach to exploit the structure of each supported input data format. For genotype formats (e.g., VCF (Danecek *et al.*, 2011)), we calculate the dot product using sparse-by-dense and sparse-by-sparse vector products. Using haplotype block format (M3VCF (Das *et al.*, 2016)), we can calculate the dot product using within-block and between-block haplotype intersections.

*Sparse Representation of Phased Genotypes* For each variant, we keep a {0, 1}^2*n*^ vector of genotypes (where 1 indicates the minor allele) and sparse vector containing the indexes of non-zero entries. If the major allele is non-reference in the input file (allele count greater than *n*), we reverse the sign of its LD coefficients for consistency. Letting *C_j_* = {*i*|*G_ij_* = 1} denote the set indexing minor-allele carriers of variant *j*, the dot product *m_jk_* := ***G**_j_* · ***G**_k_* between variants *j* and *k* can be calculated in min(*m_j_*, *m_k_*) operations, where *m_j_* is the minor allele count (MAC) for variant *j*, by using the sparse-by-dense product formula *m_jk_* = ∑_*i*∈*C_j_*_ *G_ik_*.

*Sparse Representation of Unphased Genotypes* For unphased genotypes, we store a {0, 1, 2}^*n*^ vector of genotypes and sparse vectors indexing heterozygotes and minor-allele homozygotes for each variant. In this case, LD between two variants can be calculated in min(*N*_*j*1_ + *N*_*j*2_, *N*_*k*1_ + *N*_*k*2_) operations, where *N_ji_* is the number of individuals with genotype *i* at variant *j*.

*Haplotype Block Representation* A haplotype is a sequence of contiguous alleles along a chromosome within a genomic region, or haplotype block. Due to the limited diversity of human haplotypes (Wall and Pritchard, 2003), the number of distinct haplotypes in a block with *J* biallelic variants is typically small relative to the sample size *n* or number of possible haplotypes 2^*J*^ (whichever is smaller). M3VCF format maps each sample to a haplotype within each block, and maps each variant in a block to the set of haplotypes that contain the non-reference allele (Das *et al.*, 2016). Given M3VCF input, we precompute the number of observations 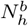 of each haplotype *h* for each block *b*, and index the set of haplotypes 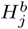 containing the minor allele at each variant *j* in block *b.* For two variants *j* and *k* in the same block, the dot product can then be calculated in at most min 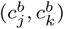 operations, where 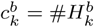 is the number of distinct haplotypes that carry the minor allele at variant *k*, using the sparse-bydense product formula 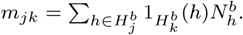 To calculate LD for variants in different blocks, we can compute a between-block count matrix 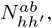 the number of samples with haplotype *h* in block *a* and haplotype *h*′ in block *b.* The dot product between variants *j* and *k* can then be calculated in 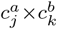 operations using the formula 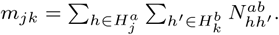 In practice, sparse-by-dense genotype products are typically more efficient for between-block calculations.

*Informed Subsampling to Estimate LD with Large Sample Sizes* When both variants *j* and *k* have large MAC (e.g., common variants and/or large sample sizes), calculating sparse-by-dense products to estimate LD becomes expensive. In this case, we use an informed subsampling approach to efficiently estimate LD while maintaining a user-specified bound on the precision of LD estimates.

We treat the sample correlation *r* = (*p_jk_*–*p_j_p_k_*)/*s_j_s_k_* as a parameter to be estimated by informed subsampling. Here, *p_j_*, *p_k_*, *s_j_* and *s_k_* can be calculated efficiently and stored; because *p_jk_* must be calculated for each pair of variants, we subsample from the carriers of the rarest allele to increase computational efficiency. In Supplementary Materials, we show that the approximate estimator *r̃_ℓ_* can be calculated in at most *ℓ* operations for any pair of variants, and increases the mean squared error (MSE) by no more than 1/*ℓ* relative to exact LD estimates (or 2/*ℓ* for unphased genotypes), where *ℓ* is a user-specified parameter. In very large data sets (*n* > 50K), subsampling with *ℓ* = 250 decreased computation time for common variants (MAF > 5%) by an order of magnitude or more.

## 3 Results

### 3.1 Implementation and Usage

We implemented our algorithms as an open-source C++ tool, emeraLD (efficient methods for estimation and random access of LD), which can be used via command line or through an R interface included with source files. emeraLD accepts block compressed VCF.gz and M3VCF.gz input, and leverages Tabix (Li, 2011) and the C library HTSlib to support rapid querying and random access of genotype data over genomic regions. emeraLD implements several options to customize output fields (variant information and LD statistics) and formats (long tables or square symmetric matrices). We also provide tools to facilitate estimating LD from a reference panel for analysis of GWAS summary statistics.

### 3.2 Performance

We used WGS genotype data from the 1000 Genomes Project Phase 3 (1KGP; *n* = 2,504) (1000 Genomes Project Consortium, 2015), Haplotype Reference Consortium (HRC; *n* = 32,470) (Haplotype Reference Consortium, 2016), and imputed genotype data from the UK Biobank (UKBB; *n* = 487,409) to compare performance between emeraLD and PLINK v1.9 (Purcell and Chang, 2016), LDstore (Benner *et al.*, 2017), VCFtools (Danecek *et al.*, 2011), and m3vcftools (Das *et al.*, 2016). For UKB, emeraLD from M3VCF.gz file input is >100× faster than PLINK from BED files (Table 1), which are >10× larger than VCF.gz and >30× larger than M3VCF.gz. For HRC, which includes 32K individuals and only variants with MAC ≥5, emeraLD calculates LD from M3VCF.gz files >6× faster than PLINK from BED files, which are >4× larger than VCF.gz and >20× larger than M3VCF.gz. Times reported for emeraLD used *ℓ* = 1, 000 (MSE of approximation ≤ 0.001); this has a negligible effect for 1KGP, but reduced overall computation time by ∼50% for UKB and HRC. Using M3VCF.gz files reduced computation time for emeraLD by ∼30-50% relative to VCF.gz.

**Table 1.** Benchmarking: Time and Memory Usage

| Tool: | m3vcftools | PLINK 1.9 | LDstore | emeraLD* | Absolute* |
| --- | --- | --- | --- | --- | --- |
| Format: | M3VCF.gz | BED | BGEN | M3VCF.gz |  |
| CPU Time Relative to emeraLD |  |  |  |  |  |
| 1KGP | 18.8 | 1.3 | 4.4 | 1.0 | 8.5 m |
| HRC | 44.7 | 6.8 | 16.8 | 1.0 | 2.6 m |
| UKB | 473.7 | 128.4 | 250.6 | 1.0 | 19.9 m |
| Memory Usage Relative to emeraLD |  |  |  |  |  |
| 1KGP | 0.7 | 137.6 | 372.4 | 1.0 | 43.8 MiB |
| HRC | 0.6 | 10.7 | 26.1 | 1.0 | 156.9 MiB |
| UKB | 0.4 | 4.7 | 4.8 | 1.0 | 4.8 GiB |

Time and memory to calculate LD in a 1Mbp region of chr20 (28,126 variants in 1KGP; 13,174 in HRC; and 32,783 in UKB). All experiments were run on a 2.8GHz Intel Xeon CPU. emeraLD \*Absolute time or memory for emeraLD as reference

### 3.3 Applications

Our approach will be implemented in a forthcoming web-based service capable of providing LD information from large panels with >60K samples, such as the TOPMed WGS project, in real time. This enables use of improved LD information by rapidly emerging and gaining in popularity web-based interactive analysis and visualization tools such as LocusZoom (Pruim *et al.*, 2010).

We have also used emeraLD to estimate LD for gene-based association and functional enrichment analysis of GWAS summary statistics.

This approach avoids precomputing and storing LD without compromising speed – for example, we developed an implementation of the MetaXcan gene-based association method (Barbeira *et al.*, 2016) using emeraLD to estimate LD on-the-fly, which is ∼5× faster than the original implementation using precomputed LD estimates. To enable simple integration with R scripts or libraries, we include an R interface to emeraLD with source files.

## 4 Conclusions

Here we described computational and statistical methods to efficiently estimate LD with large data sets. Our methods exploit two natural features of genetic data: sparsity that arises from the abundance of rare variation, and high redundancy that arises from haplotype structure. We also developed an informed subsampling approach to further improve computational efficiency while maintaining a user-specified bound on precision relative to exact LD estimates. Finally, we described an open-source software implementation that can be used to facilitate analysis of GWAS summary statistics.

## Acknowledgements

We acknowledge the developers of M3VCF and VCF format, and the cohorts and staffs of the Haplotype Reference Consortium and 1000 Genomes Project Consortium.

This research has been conducted using the UK Biobank Resource under Application Number 24460.

## Funding

The authors acknowledge support from NIH grants HG000376 (M.B.), HG007022 (G.R.A.), HG006513 (G.R.A.), and U01HL137182 (H.M.K.).

